# MAtCHap: an ultra fast algorithm for solving the single individual haplotype assembly problem

**DOI:** 10.1101/860262

**Authors:** Alberto Magi

## Abstract

**Background:** Human genomes are diploid, which means they have two homologous copies of each chromosome and the assignment of heterozygous variants to each chromosome copy, the haplotype assembly problem, is of fundamental importance for medical and population genetics.

While short reads from second generation sequencing platforms drastically limit haplotype reconstruction as the great majority of reads do not allow to link many variants together, novel long reads from third generation sequencing can span several variants along the genome allowing to infer much longer haplotype blocks.

However, the great majority of haplotype assembly algorithms, originally devised for short sequences, fail when they are applied to noisy long reads data, and although novel algorithm have been properly developed to deal with the properties of this new generation of sequences, these methods are capable to manage only datasets with limited coverages.

**Results:** To overcome the limits of currently available algorithms, I propose a novel formulation of the single individual haplotype assembly problem, based on maximum allele co-occurrence (MAC) and I develop an ultra-fast algorithm that is capable to reconstruct the haplotype structure of a diploid genome from low- and high-coverage long read datasets with high accuracy. I test my algorithm (MAtCHap) on synthetic and real PacBio and Nanopore human dataset and I compare its result with other eight state-of-the-art algorithms. All the results obtained by these analyses show that MAtCHap outperforms other methods in terms of accuracy, contiguity, completeness and computational speed.

**Availability:** MAtCHap is publicly available at https://sourceforge.net/projects/matchap/.

## Introduction

Most mammals, such as humans, are diploid organisms, which means they have two homologous copies of each chromosome, one from each parent. The two parental copies contain variants that can be homozygous (in both chromosomes) or heterozygous (in only one chromosome). While homozygous variants belong to both chromosomal copies, heterozygous alleles belong to only one of the copies and need to be partitioned into two groups.

The assignment of heterozygous variants to each chromosome copy is called phasing (or haplotype reconstruction) while the resulting sequence of variants is called haplotype and is of fundamental importance in several contexts, that include medical genetics, population genetics and association studies (Tewhey *et al*., 2011).

At present, there are two main experimental/computational approaches for phasing genomic variants: haplotype inference and haplotype assembly.

The first class of approaches comprise statistical methods (based on expectation maximization or Markov chain Monte Carlo techniques) that use genotype data collected at a set of polymorphic loci (usually SNPs) from a group of individuals and give as output the frequency of the most likely haplotype (combination of SNPs) in the population. The second class of phasing methods, haplotype assembly, aim to reconstruct the two haplotypes of a single individual by directly using sequence reads and cluster them in two groups that represent the two chromosomal copies by which they have been generated. Since 2005, the advent of second generation sequencing (SGS) has revolutionized human genomics, drastically reducing the cost of DNA sequencing and allowing the identification of single nucleotide variants, small insertions and deletions (InDels) and structural variants. However, the short length of the reads generated by SGS (up to 250 bp in paired-end sequencing) drastically limits the use these data for haplotype reconstruction, as the great majority of reads do not allow to link many (if any) variants together. As a consequence, assembled haplotypes are fragmented into short blocks that remain unphased, relative to each other.

The last few years have seen the emergence of a third generation of sequencing (TGS) technologies based on single-molecule real-time (SMRT) and nanopore sequencing, that interrogate single molecule of DNA and are capable to produce sequences in the order of tens of kb (up to hundreds of kb for nanopore) (Magi *et al*., 2016, 2017). Thanks to these new sequencing platforms, each read can span several variants along the genome allowing to infer much longer haplotype blocks.

Unfortunately, traditional haplotype assembly methods, originally devised for short sequences, fail when they are applied to TGS data, owing to the high error rate of these reads and to the fact that existing algorithmic solutions take time exponential in the number of variants per read. Although novel computational approaches, such as WhatsHap (Patterson *et al*., 2015) and HapCol (Pirola *et al*., 2016) have been developed to deal with long reads characteristics, they allow to manage only datasets with limited coverages (20x) and since the most reliable way to mitigate the high error rate of TGS reads is increasing the sequencing coverage (to 30x-50x), these novel algorithms limit the capability to reconstruct haplotypes of higher quality.

To overcome the limits of currently available approaches, I introduce a novel formulation of the objective function for solving the single individual haplotype assembly problem, based on maximum allele co-occurrence (MAC) and I propose a very fast algorithm that allows to analyze high coverage TGS datasets with times comparable to low coverage ones. I test my algorithm on simulated and real sequencing data and I show that it outperforms other state of the art methods in terms of accuracy, contiguity, completeness and computational speed.

## Materials and Methods

### Maximum allele co-occurrence (MAC) function

In the last decade several computational strategies have been developed to reconstruct the two haplotypes of a single individual by directly using sequence reads (single individual haplotype assembly problem) and are all based on the fragment matrix that is defined as an *n* × *m* matrix where *n* in the number of heterozygous variants and and *m* is the number of reads (fragments).

Three main strategies have been used to solve the single individual haplotype assembly problem: removal of fragment matrix rows (Minimum Fragment Removal, MFR), removal of fragment matrix columns (Minimum SNP Removal, MSR) and flipping of fragment matrix elements (Minimum Error Correction, MEC). All these problems are NP-hard (Cilibrasi *et al*., 2007) and the approach that has received most attention in literature is the MEC problem that aims to find the minimum number of error corrections in order to partition the reads in two haplotypes.

Many algorithms have been proposed to solve the MEC problem and these include exact methods based on Integer Linear Programming and Dynamic Programming (Chen *et al*., 2013; He *et al*., 2010) and heuristic approaches (Panconesi *et al*., 2004; Duitama *et al*., 2012; Edge *et al*., 2017; Bansal *et al*., 2008). These methods obtain good results with short read data, but fail when applied to TGS data since they scale poorly with increasing read length.

On the other hand, recently developed algorithms, such as WhatsHap (Patterson *et al*., 2015) and HapCol (Pirola *et al*., 2016), properly devised for long TGS reads, can deal only with datasets of limited coverages (20x).

In order to overcome the limits of currently available algorithms, I propose a novel formulation of the haplotype assembly problem that aims to infer the two haplotypes that maximize the number of allele co-occurrence in the input fragments.

In a sequencing experiment, each read represents a fragment of a chromosome and when it spans more than a heterozygous variant it represents a segment of one of the two haplotypes. In this situation, the co-occurence of two alleles in the same read supports the fact that one haplotype contains the two alleles, while the other haplotype contains the two other alleles. As a consequence, one can argue that the higher the number of reads containing two alleles and the higher the probability the two alleles belong to the same haplotype.

From this considerations, one can infer the two haplotypes of a diploid genome by maximizing the allele co-occurrence objective function (MAC) that represent the total number of allele co-occurence in the fragment matrix:

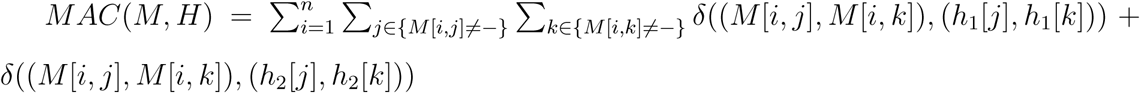

where *δ*((*a, b*), (*c, d*)) is a function equal to 1 if *a* = *c* and *b* = *d*, equal to 0 elsewhere, while M, the fragment matrix, represent the input to *MAC* function that consists of *n* rows (reads) and *m* columns (heterozygous variants).

Each row of M represents the information of a read and each column the information of a heterozygous variant. The information is encoded as a string that takes values on the alphabet 0, 1, −. *M* [*i, j*] = − means that fragment *i* does not cover variant *j, M* [*i, j*] = 0 (*M* [*i, j*] = 0) means that fragment *i* has reference (alternative) allele at locus *j. h*_1_ and *h*_2_ are the two solution haplotypes (vectors of length N where N is the number of heterozygous variants), with *h* ∈ {0, 1} and since each variant is heterozygous, *h*_1_[*j*] ≠ *h*_2_[*j*] for each *j*.

In order to maximize the MAC objective function I developed an algorithmic recipe that recursively infer the values of *h*[*j*] = {*h*_1_[*j*], *h*_2_[*j*]} by maximizing the variant specific MAC function:

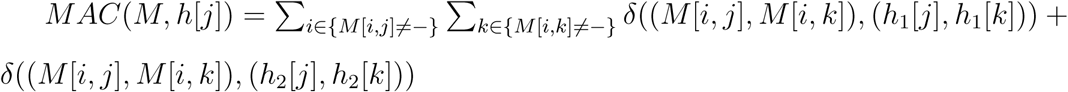

that represent the total number of allele co-occurence for variant *j*.

The pseudo code of the Maximum Allele Cooccurrence Haplotype (MAtCHap) algorithm is the following:

1. randomly initialize *h*^1^ so that *h*^1^[1] = {1, 0} (or *h*^1^[1] = {0, 1}), in this way *h*^1^ = ({0, 1}, {−, −}….{−, −})
  2. repeat until *h*^*t*^ ≠ *h*^*t*−1^
  3. for j=2 to m
    4. Calculate *MAC*(*M, h*^*t*^[*j*]) for *h*^*t*^[*j*] = {0, 1} and *h*^*t*^[*j*] = {1, 0} and assign *h*^*t*^[*j*] the value that give *max*(*MAC*(*M, h*^*t*^[*j*]))
  5. t=t+1

The solution that one obtains from this procedure is the two haplotypes *h*^*t*^ that maximize the number of allele co-occurrence in the fragment matrix *M* (Figure 1). At the end of the maximization procedure, for each variant *j* we calculate the ratio 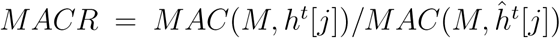 (where 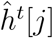 is the flipped version of *h*^*t*^[*j*] with inverted alleles) that gives a measure of the likelihood that the two alleles where correctly assigned to their haplotypes. *MACR* close to 1 means that the number of allele co-occurrence that support *h*^*t*^[*j*] and 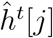 are very similar, while high values of *MACR* mean that the number of allele co-occurrence that support *h*^*t*^[*j*] are much larger than those supporting 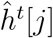 and so the likelihood of *h*^*t*^[*j*] is much higher than 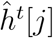. Positions with low *MACR* can be filtered out to improve the accuracy of haplotype assembly. Owing to low-coverage and highly error prone genomic positions, haplotype inference can produce switch errors that lead to completely flipped haplotype blocks.

**Figure 1:**
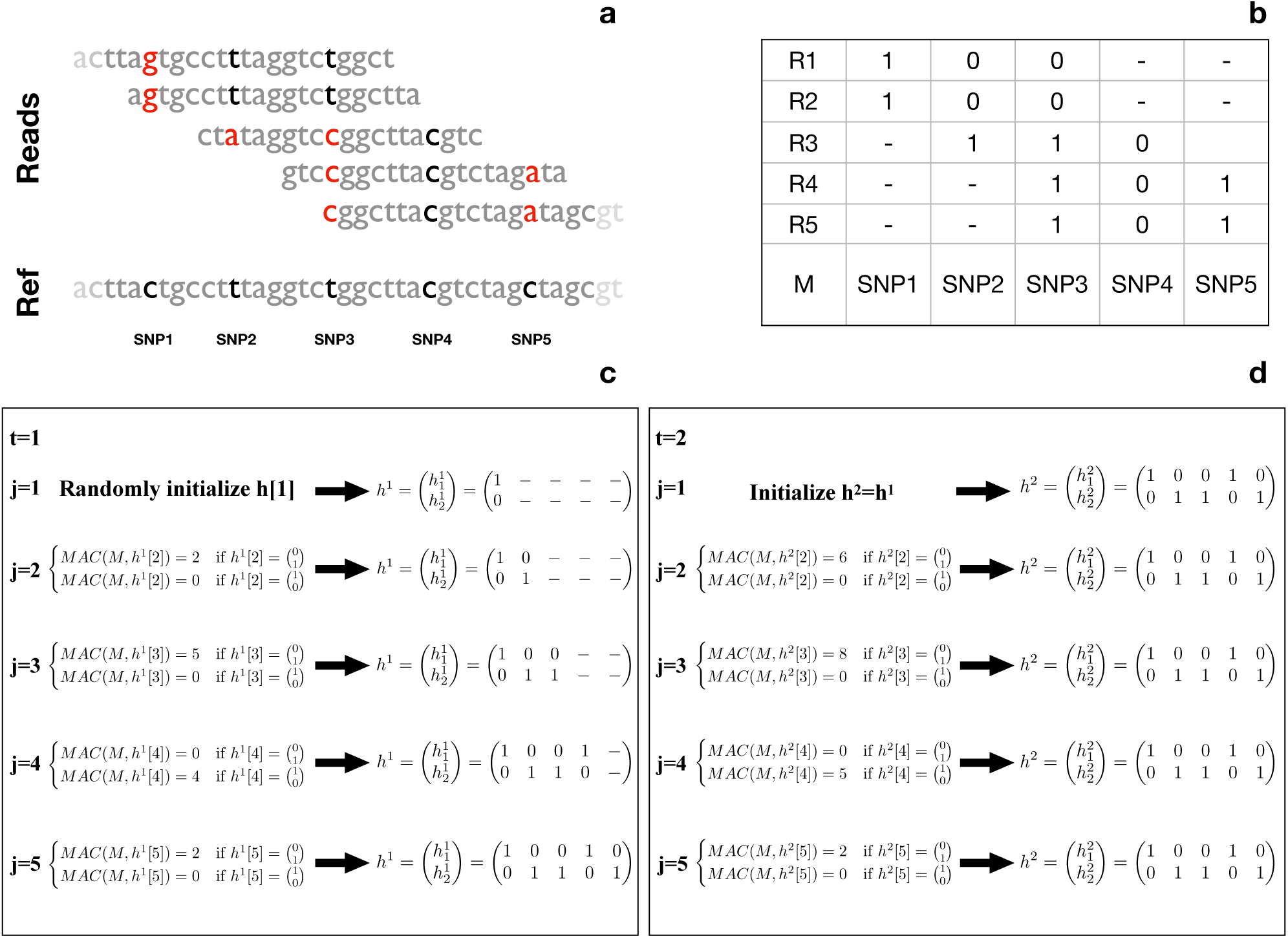
MAtCHap algorithm scheme. In panel (a) is reported the alignment of five reads against a segment of the reference genome that spans five heterozygous variants. Panel (b) reports the fragment matrix M for the five reads. Panels (c) and (d) show the MAtCHap algorithm applied to matrix M for *t* = 1 (c) and *t* = 2 (d).

To overcome this problem, at the end of the maximization procedure, the MAtCHap algorithm include a procedure to remove switch errors by calculating partial MAC function:

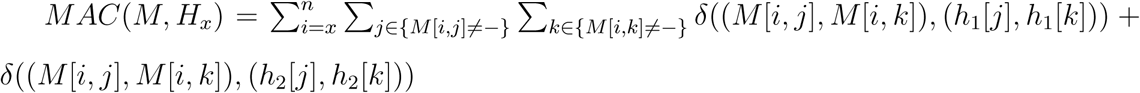

Partial MAC function *MAC*(*M, H*_*x*_) smaller than 1 indicate a switch error at position *x* and every position from *x* onwards is flipped with respect to those before *x*.

### Synthetic data simulations

To assess the performance of MAtCHap algorithm in reconstructing individual haplotypes from long and highly error prone reads, synthetic sequences were generated by using the PBSim software (Ono *et al*., 2013). PBSim simulates reads by randomly sampling from a reference sequence, adding errors with a user defined distribution of substitutions, insertions and deletions and allowing to define read size distribution (mean and maximum size) and the desired sequencing coverage.

In all the performed simulations, the two haplotypes were generated by PBSim using as reference sequences two modified versions of 10 Mb of the human reference genome GRCh37 (chr1:10.000.001-20.000.000), in which synthetic substitutions were added every N bp (for each haplotype *N* was set to *N* = {200*bp*, 1000*bp*, 2000*bp*, 4000*bp*} obtaining a diplotype with a heterozygous variant every 100 bp, 500 bp, 1000 bp and 2000 bp respectively).

### Real data

The nanopore WGS consortium (Jain *et al*., 2018) (https://github.com/nanopore-wgs-consortium) sequenced the NA12878 human genome on the ONT MinION by using 39 R9/R9.4 flow cells generating 4,183,584 base-called reads containing 91,240,120,433 bases with a read N50 of 10,589 bp. Reads in FastQ format for all 39 runs were downloaded from https://github.com/nanopore-wgs-consortium, aligned against the human reference genome (hg19) by using BWA mem (Li *et al*., 2010) with −*xont*2*d* option and converted to bam format with samtools (Li *et al*., 2009) obtaining a 30x sequencing coverage. Bam file was then downsampled (with samtools -s) to simulate datasets with sequencing coverage of 5x, 10x, 15x, 20x, 25x and 30x.

The Genome in a Bottle (GIAB) Consortium (https://github.com/genome-in-a-bottle) is creating diverse set of sequencing data for seven human genomes that include the NA12878 from the CEPH Utah Reference collection (Zook *et al*., 2016). The sequencing data were obtained from 11 technologies: BioNano Genomics, Complete Genomics paired-end and LFR, Ion Proton exome, Oxford Nanopore, Pacific Biosciences, SOLiD, 10X Genomics GemCodeTM WGS, and Illumina paired-end, mate-pair, and synthetic long reads.

NA12878 genome was sequenced with the Pacific Biosciences Single Molecule Real-Time (SMRT) platform using the P6-C4 chemistry and obtaining a 45x total sequencing coverage. The bam file of PacBio reads mapped to the human reference genome (hg19) with BLASR (v1.3.2) were downloaded from ftp://ftp-trace.ncbi.nih.gov/giab/ftp/data/NA12878/NA12878_PacBio_MtSinai/. To simulate different sequencing coverage (5x, 10x, 15x, 20x, 25x and 30x), the original 45x bam file was downsampled with samtools view -s.

To evaluate the performance of MAtCHap and the other eight state-of-the-art algorithms we used as gold standard the high confidence variants identified by the GIAB consortium and downloaded at ftp://ftp-trace.ncbi.nlm.nih.gov/giab/ftp/release/ NA12878_HG001/latest/GRCh37/. VCFs include phasing information both from family-based phasing and from local read-based phasing, prioritizing family-based phasing. 99.0% of high-confidence calls are phased by the Platinum Genome (Eberle *et al*., 2017).

### Single Individual Haplotype Tools

The performance of MAtCHap were compared with those obtained by other height state of the art tools: WhatsHap (Patterson *et al*., 2015), HAPCUT2 (Edge *et al*., 2017), TwoDMEC (Wang *et al*., 2007), FastHare (Panconesi *et al*., 2004), DGS (Levy *et al*., 2007), ProbHap (Kuleshov *et al*., 2014), MixSIH (Matsumoto *et al*., 2013) and RefHap (Duitama *et al*., 2012).

WhatsHap v0.18 was downloaded at https://whatshap.readthedocs.io/en/latest/ and applied to synthetic and real datasets by following the recommendations reported in https://whatshap.readthedocs.io/en/latest/guide.html#recommended-workflow. HAPCUT2 https://github.com/vibansal/HapCUT2 was applied to PacBio and Nanopore synthetic and real datasets with default parameter settings. TwoDMEC, FastHare, DGS and Refhap algorithms were ran exploiting the SingleIndividualHaplotyper package downloaded at http://www.molgen.mpg.de/~genetic-variation/SIH/Data/algorithms. Prob-Hap was downloaded at https://github.com/kuleshov/ProbHap and ran following author suggestions.

MixSIH https://github.com/hmatsu1226/MixSIH was ran by setting the error rate option (-a) to 0.15 for real Nanopore and PacBio data (according to nanopore and PacBio error profiles) and to 1 − *error* for synthetic data (where error is the global error rate simulated by PBSim).

## Results

### Synthetic Simulations

To demonstrate the ability of MAtCHap to reconstruct haplotypes from sequences with different sizes and error rates, the PBSim software was used to simulate synthetic data (see methods).

In particular, to test the capability of my computational recipe to infer the correct haplotypes from sequences generated by novel TGS platforms, PBSim was exploited to simulate sequencing dataset that mimic the characteristics of long reads with average size from 5 Kb to 15 Kb (size={5 Kb, 8 Kb, 10 Kb, 12 Kb and 15 Kb}), global error rate from 75% to 99% (error={0.75,0.8, 0.85, 0.9, 0.95, 0.99}) and total sequencing coverage from 5x to 50x (coverage={5x,10x, 15x, 20x, 25x, 30x, 35x, 40x, 45x and 50x}). Moreover, to better simulate the sequences produced by PacBio and MinION platforms, the error rate distribution among substitutions, insertions and deletions, were set according to the results found by me and other authors (Magi *et al*., 2016; Weirather *et al*., 2017): 10:60:30 for PacBio and 30:30:40 for MinION.

Assessing the quality of an haplotype assembly requires very complex measures that take into consideration accuracy, contiguity and completeness of the inferred haplotype. In an inferred haplotype one can find two type of errors: switch and mismatch errors. Switch errors (also known as long switch) are defined as two or more consecutive variants flipped with respect to gold standard, while mismatch errors are defined as variants whose phase was incorrectly inferred (also known as short switch). A switch error (also known as long switch) occurs when the phase between two adjacent variants in the assembled haplotypes is discordant relative to the truth haplotypes. Two consecutive switch errors correspond to the flipping of the phase of a single variant and were counted as ‘mismatch’ (also known as short switch) errors instead of two switch errors.

The accuracy of a haplotype assembly is typically evaluated by comparing the inferred haplotypes to gold standard haplotypes and calculating the total fraction of switch and mismatch errors (Edge *et al*., 2017).

While total error rate (switch + mismatch) gives a global picture of haplotype accuracy, it is not capable to capture the effect of short- and long-range switch errors on the reconstructed haplotype structure. The limits of total error rate can be overcome by using pairwise variant haplotype assignment accuracy, in which inferred alleles at any given pair of positions are compared to gold standard calls and classified as concordant or discordant. Comparing every pair of variants, binned by genomic distances, one can study the effect short- and long-range switch errors on haplotype accuracy.

Concerning the evaluation of haplotype contiguity (how many separate haplotype fragments have been inferred), a standard measure is the N50 metric, that was originally developed for the comparative assessment of de novo genome assemblies, and that is defined as the length of the shortest haplotype at 50% of the total assembly length. However, since this statistic does not take into account accuracy, long and highly erroneous blocks can obtain N50 measures higher than short and accurate haplotypes. One way to resolve this limitation is to use a contiguity measures that breaks haplotype blocks at each erroneous position (switch and mismatch error) and then calculating the N50 of the resulting assembly.

Moreover, also statistics that evaluate the proportion of all heterozygous variants that have been phased (completeness) are highly informative for assessing the quality of inferred haplotypes.

For these reasons, for evaluating the performance of the MatchHap algorithm I used four different measures that capture accuracy, contiguity and completeness and I compared its performance with those obtained by other eight state-of-the-art methods: WhatsHap, HAPCUT2, TwoDMEC, FastHare, DGS, ProbHap, MixSIH and RefHap (see Methods). MatchHap is one of the few algorithms which can provide estimates of haplotype membership of a variant in the form of *MACR* score that a heterozygous variant belong to the assigned haplotype. To evaluate the capability of the *MACR* score to predict haplotype reconstruction accuracy, all the variants were first ranked as a function of *MACR* score and then the total fraction of switch and mismatch errors was calculated by considering only variants above a given score threshold (and treating the other variants as unphased). These results were compared with the only other tools that allows the user to exclude low-confidence positions (HAPCUT2, ProbHap and MixSIH) and they show (Figure 2) that MAtCHap obtains the lowest error rate for each fraction of phased SNVs at each sequencing coverages, followed by MixSIH, HAPCUT2 and ProbHap respectively.

**Figure 2:**
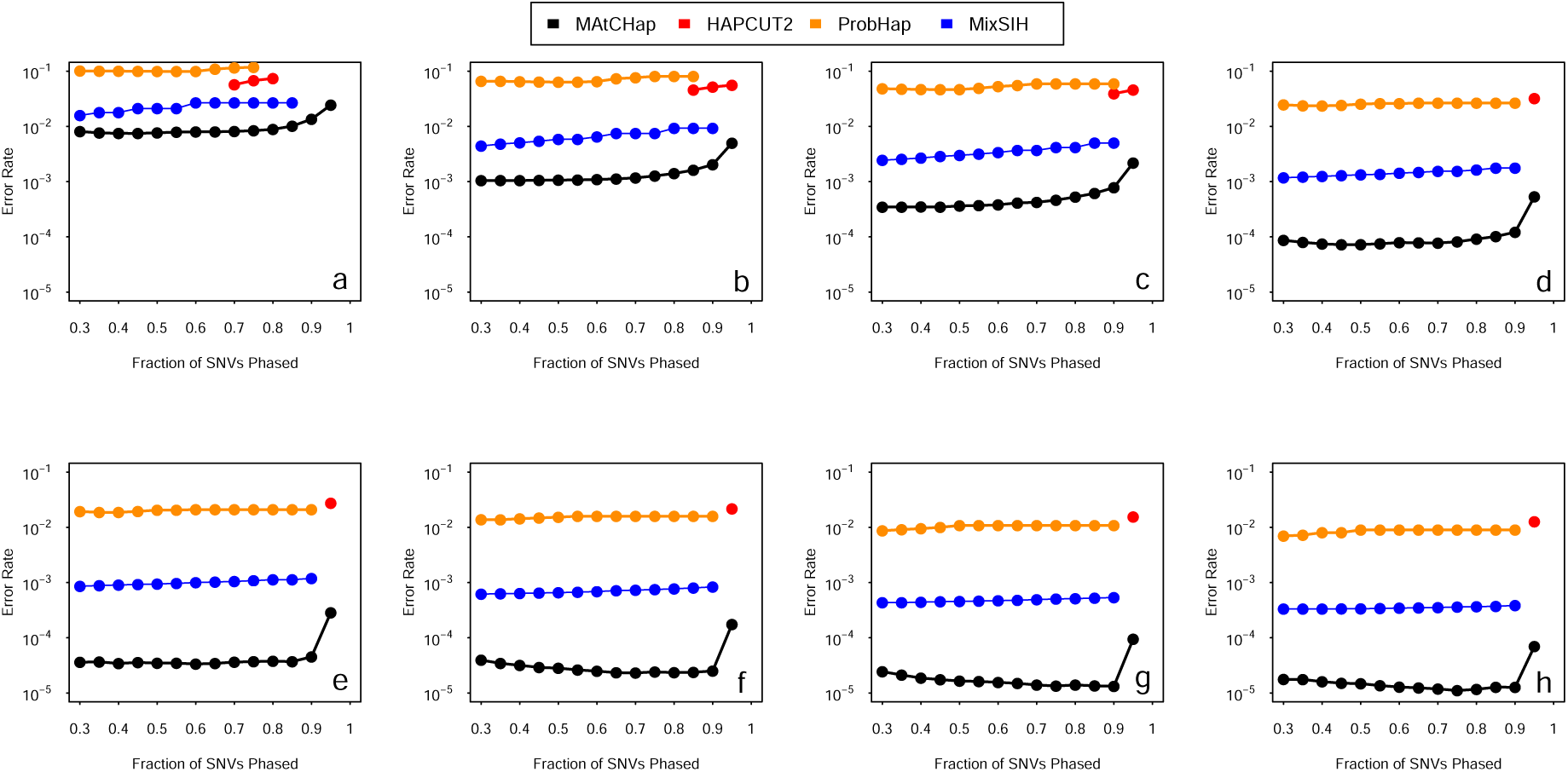
Comparison of the error rate/completeness trade-off of MAtCHap, HAPCUT2, PROBHAP and MixSIH as a function of haplotype score. For each of the four tools, all the variants were ranked as a function of tool specific haplotype score and, by considering only variants above a given score threshold, we then calculated error rate and percentage of phased variants. Panels report the results obtained by the four tools for different sequencing coverages: 5x (a), 10x (b), 15x (c), 20x (d), 25x (e), 30x (f), 40x (g), 50x (h).

As a further step, to evaluate the global error rate of each algorithm I calculated the sum of mismatch and switch errors and to capture the effect of short and long-range errors, I estimated pairwise accuracy as a function of distance by calculating the proportions of concordant alleles at any pairs of variants using bin of distances that range from 100 to 900 Kb (see methods). The results of Figure 3.a-f show that MAtCHap is capable to better reconstruct both the short- and long-range structure of haplotypes (panels a and b) generating the lowest global error rate for each sequencing coverage and accuracy, followed by RefHap and MixSIH. Surprisingly, two recently published algorithms, WhatsHap and HAPCUT2, properly devised for haplotype assembly from long reads, obtained poor performance.

**Figure 3:**
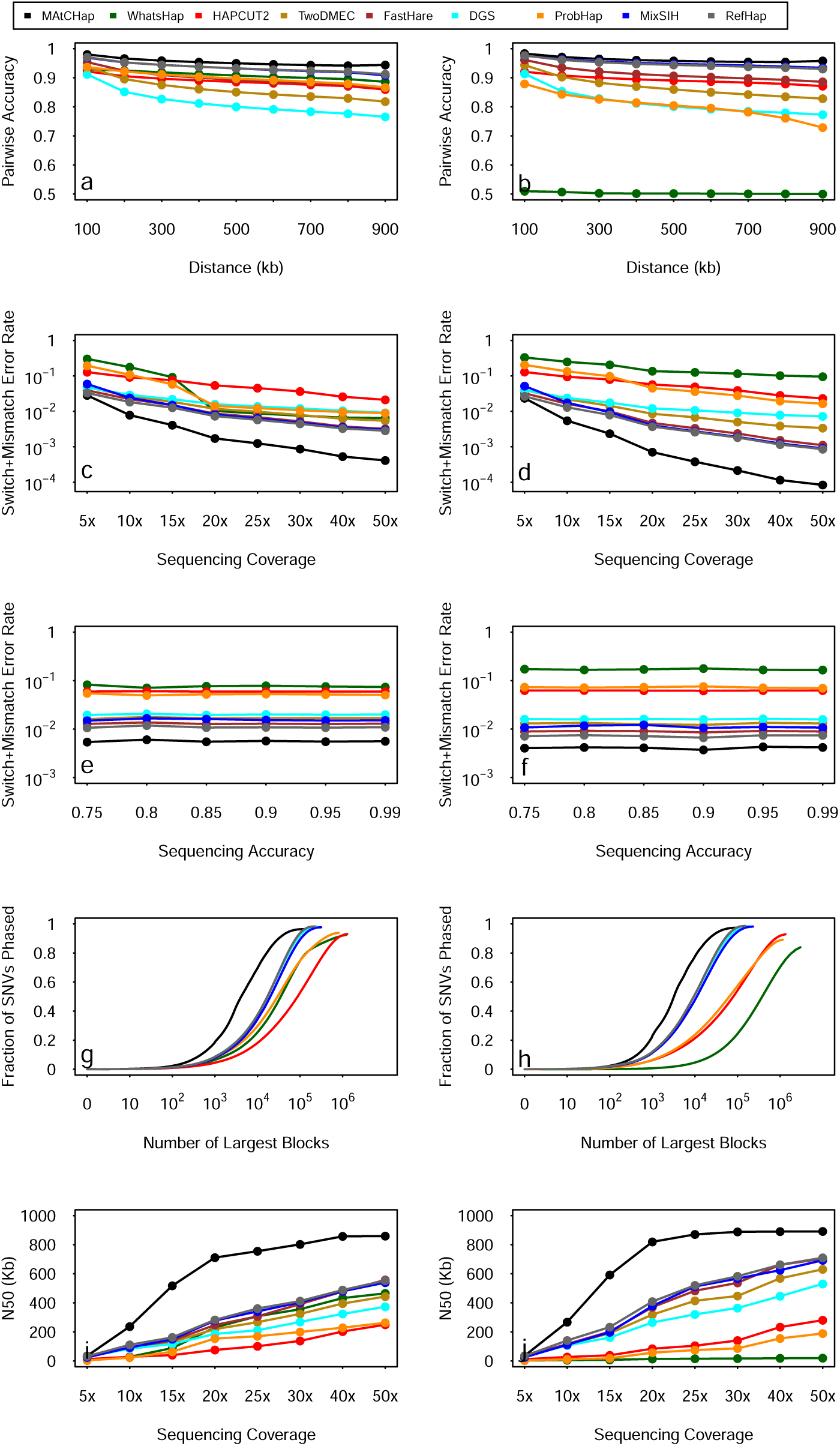
Accuracy, completeness and contiguity of haplotype assembly for simulated data. Panels a-b report pairwise accuracy as a function of genomic distance. Panels c-f report the total error rate (switch + mismatch errors) as a function of sequencing coverage (c-d) and read accuracy (e-f). Panels g-h show haplotype completeness as fraction of phased variants vs number of inferred blocks. N50 contiguity measure vs sequencing coverage is reported in panels i-j. Panels a, c, e, g and i report the results for simulated Nanopore reads, while panels b, d, f, h and j for simulated PacBio reads.

Finally, to evaluate contiguity and completeness of the inferred haplotypes, N50 (with haplotype blocks broken at each errors) and the proportion of phased variants as a function of the number of largest blocks were calculated. Panels g-j of Figure 3 show that my approach outperforms other algorithms in term of completeness (Figure 3.g-h) and that is capable to obtain high N50 measures even with moderately low coverages (20x) while the other tools requires sequencing coverages as large as 50x.

### Real data analysis

To assess the performance of the MAtCHap algorithm to infer haplotypes from real data I exploited two TGS dataset generated with Nanopore and PacBio technologies on the NA12878 sample.

The Genome in a Bottle (GIAB) Consortium (https://github.com/genome-in-a-bottle) sequenced NA12878 with the Pacific Biosciences Single Molecule Real-Time (SMRT) platform using the P6-C4 chemistry and obtaining a 45x total sequencing coverage (see Methods for more details). Similarly, the nanopore WGS consortium (https://github.com/nanopore-wgs-consortium) sequenced the NA12878 human genome with the ONT MinION by using 39 R9/R9.4 flow cells generating 30x total sequencing coverage dataset (see Methods for more details).

In order to evaluate the capability of my algorithm to reconstruct the haplotype structure of a human genome with data at different sequencing coverage, the original TGS datasets were downsampled to simulate 5x, 10x, 15x, 20x, 25x and 30x. As in Synthetic Simulations section, MAtCHap and the other eight state-of-the-art tools were evaluated with four different statistics that measure accuracy, contiguity and completeness by using as gold standard the variants phased by the GIAB consortium (see Methods for more details).

The results reported in Figure 4 clearly show that my approach gives the best results in terms of accuracy, obtaining the lowest error rate and the highest pairwise accuracy for both PacBio and Nanopore data at all the sequencing coverages, followed by HAPCUT2 and WhatsHap.

**Figure 4:**
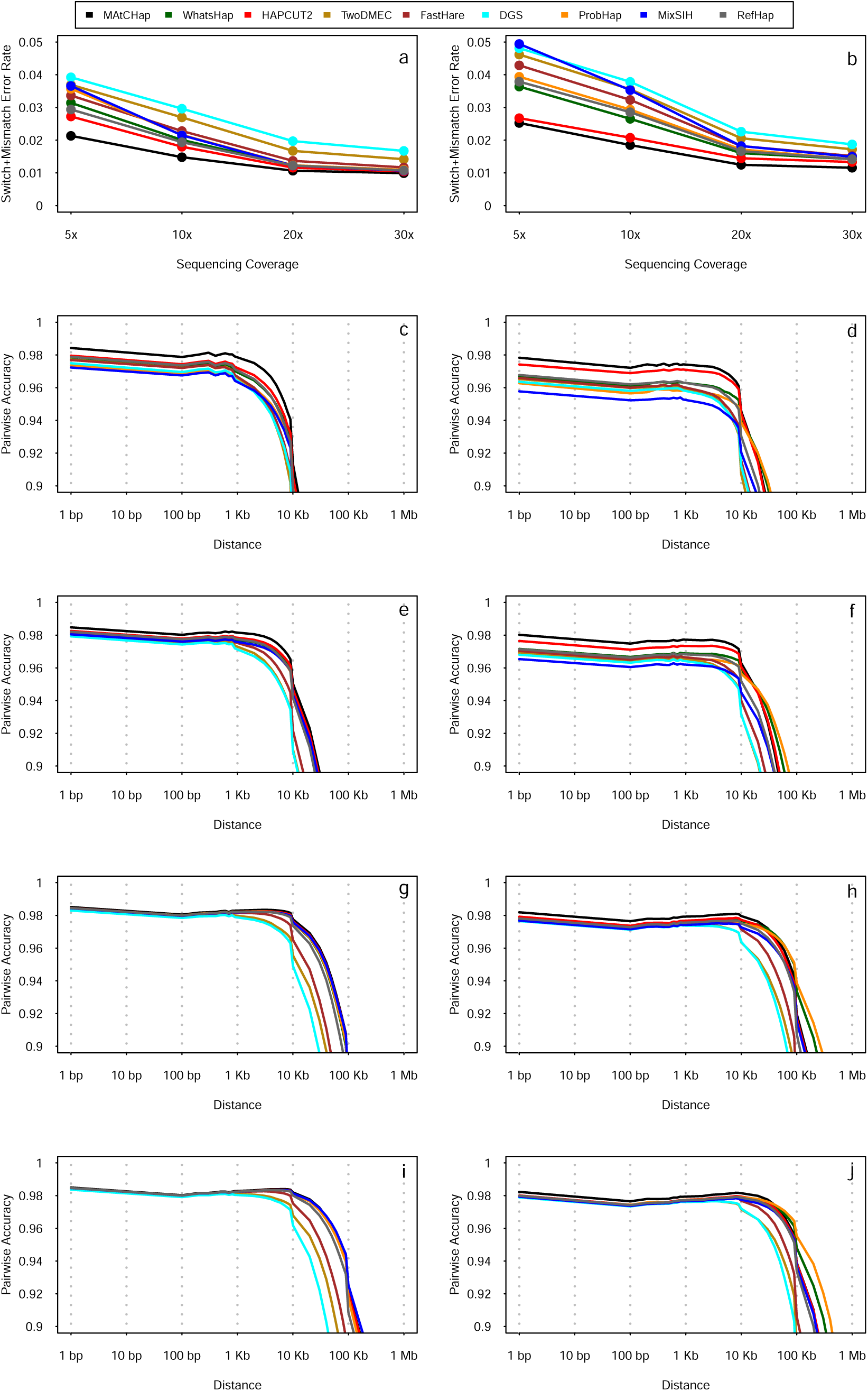
Accuracy plot for NA12878 Nanopore and PacBio sequencing data. Panels a-b show the total error rate (switch + mismatch errors) as a function sequencing coverage, while panels c-j report pairwise accuracy as a function of genomic distance (c-d for 5x, e-f for 10x, g-h for 20x and i-j for 30x). Panels a, c, e, g and i report the results for PacBio experiments and panels b, d, f, h and j for Nanopore.

Concerning completeness and contiguity, Figure 5 show that although all the algorithms obtain very similar results, my method outperforms the other tools in terms of both fraction of phased variants and N50 size for PacBio and Nanopore data. In contrast with the results obtained with simulated data, HAPCUT2 and WhatsHap tools obtained good results in the analyses of real TGS dataset, demonstrating their capability to manage the intrinsic characteristics of this new generation of sequencing data.

**Figure 5:**
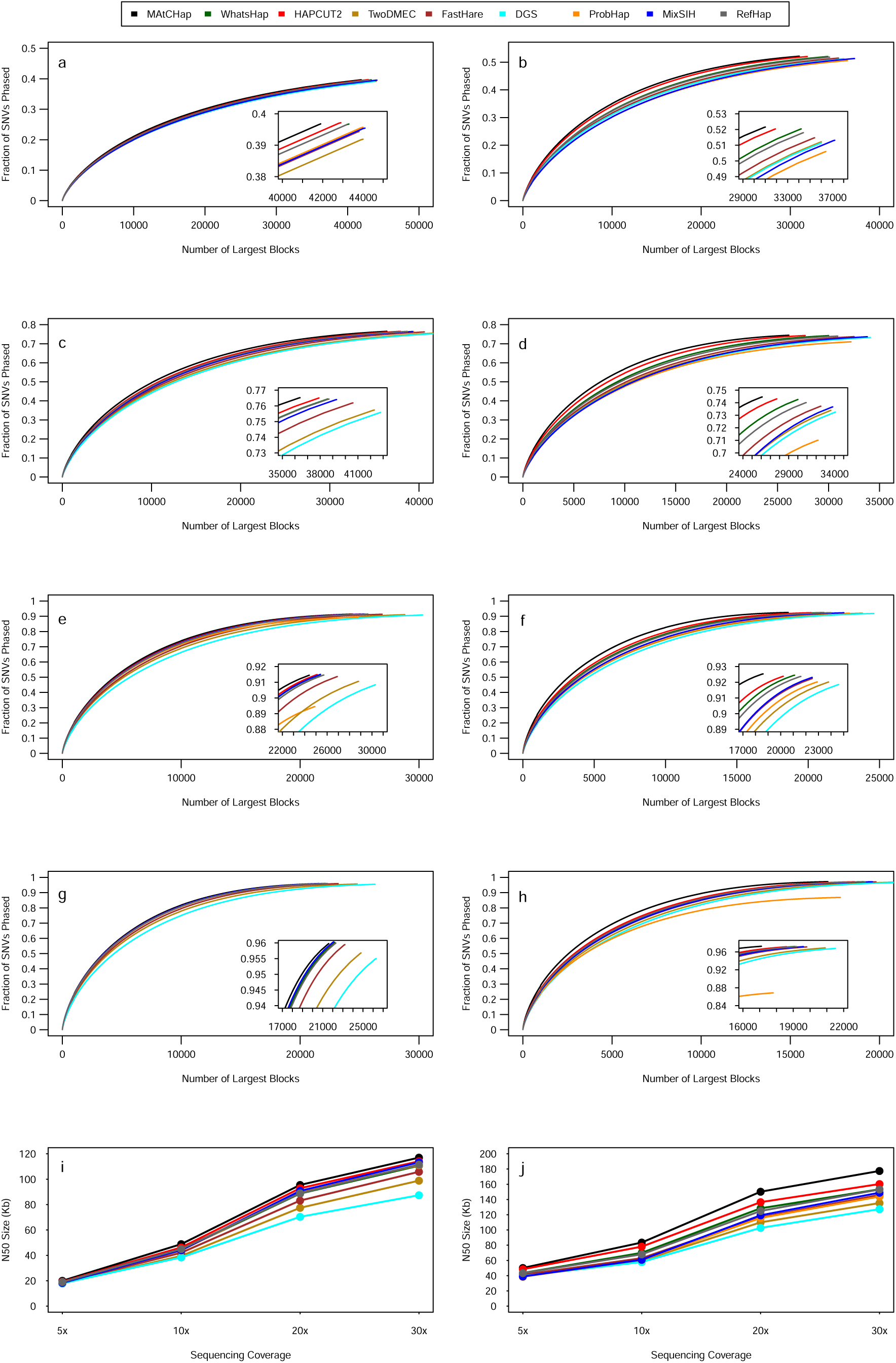
Completeness and contiguity plots for NA12878 Nanopore and PacBio sequencing data. Panels a-h show haplotype completeness as fraction of phased variants vs number of inferred blocks for sequencing coverage at 5x (a-b), 10x (c-d), 20x (e-f) and 30x (g-h). Panels i-j show the N50 contiguity measure as a function of sequencing coverage. Panels a, c, e, g and i report the results for PacBio experiments and panels b, d, f, h and j for Nanopore.

Finally, in order to show the computational performance of my method, I calculated the time taken by the nine algorithms to infer the haplotypes of all the 22 autosomal chromosomes for TGS data at different sequencing coverages. As expected, the results of figure 6 show that while the computational performance of all MEC-based methods are highly influenced by total number of reads, MAtCHap obtains nearly the same times at each sequencing coverage, demonstrating its power and versatility in the analysis of future ultra high throughput long reads datasets.

**Figure 6:**
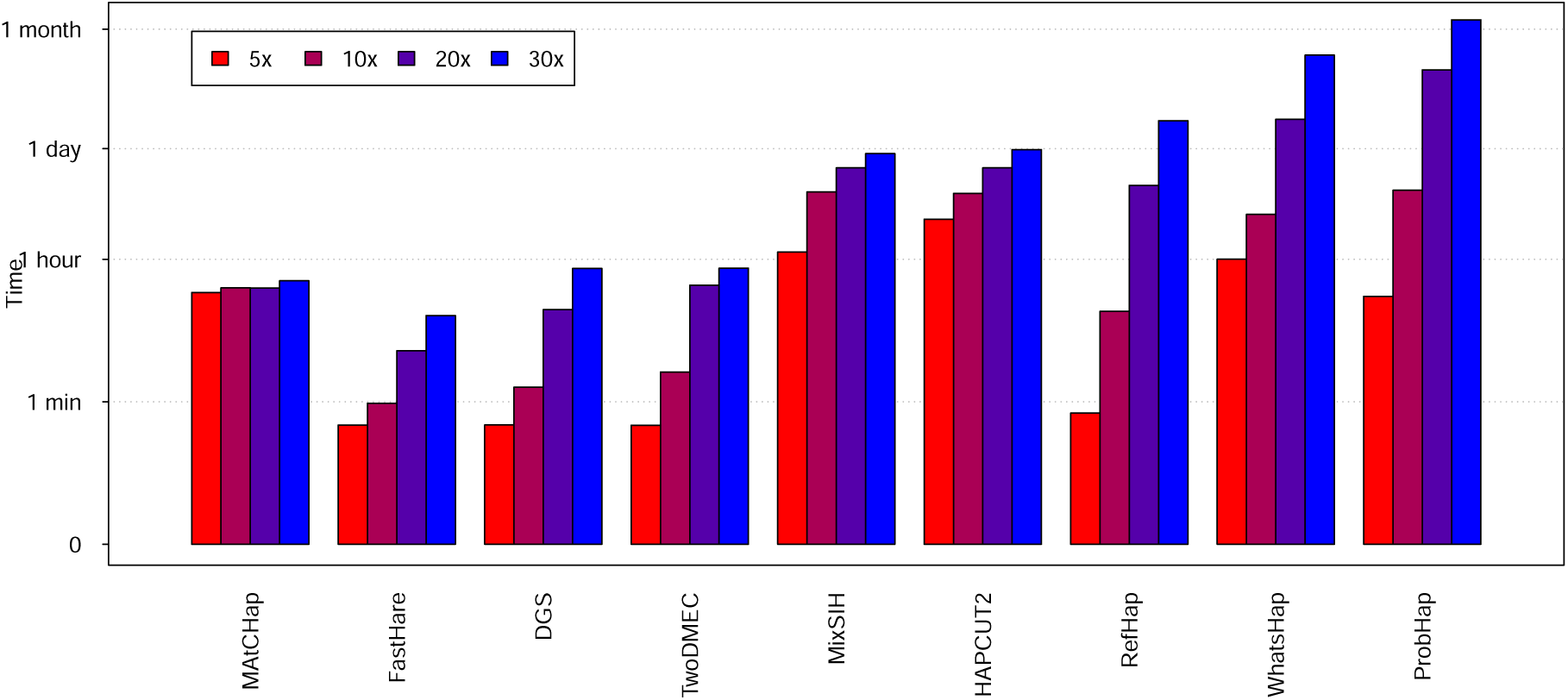
Computational performance of nine algorithms in the analysis of real long read data. The barplots report the computational time taken by the nine haplotype assembly algorithms to infer the haplotypic structure of the NA12878 genome with long reads at 5x, 10x, 20x and 30x of sequencing coverage.

## Discussion and Conclusion

From the advent of second generation sequencing technologies, the vast majority of genomic studies have ignored the diploid nature of the human genome owing to the computational complexity and the lack of cost-effective experimental approaches for obtaining phase information. However, many evidences, coming from classical medical genetics literature and from recent studies, demonstrate that the correlation between genotype and phenotype can be more fully understood by using phase information (Tewhey *et al*., 2011). In medical genetics applications, haplotype information can be of fundamental importance in the study of compound heterozygous variants that occur when two homologous copies of a gene contain unique variants at different positions. These variants can perturb the function of the two homologous copies of a gene, generating a phenotype that is different from that seen if one homologous gene carries both deleterious variants.

Compound heterozygosity is the genetic mechanism at the base of several recessive Mendelian disorders such as Hemachromatosis, Mediterranean Fever and many other. Compound heterozygosity is also at the base of the “two hit” model of cancer, in which an individual with an inherited cancer-susceptibility variant develops a somatic mutation at a different position in the same gene that leads to dysfunction in both gene copies and to potential tumorigenic effect. In compound heterozygous studies, genotype information at different loci is not sufficient, while is essential the knowledge of haplotype.

In this scenario, the development of low-cost and highly efficient experimental/computational strategies for the reconstruction of haplotype structure is of fundamental importance for human genomics. Thanks to the advent of third generation nanopore sequencing, in particular the novel Promethion platform, in the next few years it will be possible to sequence a high coverage human genome (30-40x, and even 50x) at affordable prices comparable to Illumina short reads technology. For these reasons, the availability of powerful and fast algorithms that are capable to handle this new generation of sequencing data without being influenced by the total throughput is of fundamental importance.

In this work I introduce a novel formulation of the single individual haplotype assembly problem based on the concept of maximum allelle co-occurrence (MAC), I define an objective function that mathematically represent the MAC formulation and I propose a fast algorithmic recipe (MAtCHap) that maximizing the MAC function is capable to reconstruct the haplotype structure of a diploid genome from sequencing data.

To show the power of my computational approach in the analysis of novel TGS long read datasets, MAtCHap was tested on simulated and real PacBio and Nanopore experiments and was compared with other eight state-of-the-art methods based on classical MEC formulation of the haplotype assembly problem. To evaluate the performance of the nine algorithms I used several statistics that measure different properties of the reconstructed haplotypes, and in all the analyses I performed, my method outperformed the other height algorithms in terms of accuracy, contiguity and completeness.

Moreover, MAtCHap obtained the best performance also in terms of computational performance, allowing to infer high quality haplotypes from a 30x sequencing coverage long read experiment in less than one hour, while other tools can take from one day (HAPCUT2) to 1 month (WhatsHap) with lower accuracy. Remarkably, while the computational performance of other tools are highly influenced by sequencing coverage, MAtCHap is capable to analyze a 30x experiment with times comparable to a 5x experiment, demonstrating high scalability for future ultra-high coverage TGS experiments.

## References

Bansal V, Bafna V. HapCUT: an efficient and accurate algorithm for the haplotype assembly problem. Bioinformatics. 2008 Aug 15;24(16):i153–9.

Chen ZZ, Deng F, Wang L. Exact algorithms for haplotype assembly from whole-genome sequence data. Bioinformatics. 2013 Aug 15;29(16):1938–45.

Cilibrasi R, Van Iersel L, Kelk S, Tromp J. The complexity of the single individual SNP haplotyping problem. Algorithmica. 2007; 49(1):13?36.

Duitama J, McEwen GK, Huebsch T, Palczewski S, Schulz S, Verstrepen K, Suk EK, Hoehe MR. 2012. Fosmid-based whole genome haplotyping of a HapMap trio child: evaluation of single individual haplotyping techniques. Nucleic Acids Res 40: 2041–2053.

Eberle MA, Fritzilas E, Krusche P, Kllberg M, Moore BL, Bekritsky MA, Iqbal Z, Chuang HY, Humphray SJ, Halpern AL, Kruglyak S, Margulies EH, McVean G, Bentley DR. A reference data set of 5.4 million phased human variants validated by genetic inheritance from sequencing a three-generation 17-member pedigree. Genome Res. 2017 Jan;27(1):157–164.

Edge P, Bafna V, Bansal V. HapCUT2: robust and accurate haplotype assembly for diverse sequencing technologies. Genome Res. 2017 May;27(5):801–812.

He D, Choi A, Pipatsrisawat K, Darwiche A, Eskin E. Optimal algorithms for haplotype assembly from whole-genome sequence data. Bioinformatics. 2010 Jun 15;26(12):i183–90.

Jain M, Koren S, Miga KH, Quick J, Rand AC, Sasani TA, Tyson JR, Beggs AD, Dilthey AT, Fiddes IT, Malla S, Marriott H, Nieto T, O’Grady J, Olsen HE, Pedersen BS, Rhie A, Richardson H, Quinlan AR, Snutch TP, Tee L, Paten B, Phillippy AM, Simpson JT, Loman NJ, Loose M. Nanopore sequencing and assembly of a human genome with ultra-long reads. Nat Biotechnol. 2018 Apr;36(4):338–345.

Kuleshov V. Probabilistic single-individual haplotyping. Bioinformatics. 2014 Sep 1;30(17):i379–85.

Levy S, Sutton G, Ng P, Feuk L, Halpern A, Walenz B, Axelrod N, Huang J, Kirkness E, Denisov G, et al. 2007. The diploid genome sequence of an individual human. PLoS Biol 5: e254.

Li H, Handsaker B, Wysoker A, Fennell T, Ruan J, Homer N, Marth G, Abecasis G, Durbin R; 1000 Genome Project Data Processing Subgroup. The Sequence Alignment/Map format and SAMtools. Bioinformatics. 2009 Aug 15;25(16):2078–9.

Li H, Durbin R. Fast and accurate long-read alignment with Burrows-Wheeler transform. Bioinformatics. 2010 Mar 1;26(5):589–95.

Magi A, Giusti B, Tattini L. Characterization of MinION nanopore data for resequencing analyses. Brief Bioinform. 2017 Nov 1;18(6):940–953.

Magi A, Semeraro R, Mingrino A, Giusti B, D’Aurizio R. Nanopore sequencing data analysis: state of the art, applications and challenges. Brief Bioinform. 2018 Nov 27;19(6):1256–1272.

Matsumoto H, Kiryu H. 2013. MixSIH: a mixture model for single individual haplotyping. BMC Genomics 14: S5.

Ono Y, Asai K, Hamada M. PBSIM: PacBio reads simulator–toward accurate genome assembly. Bioinformatics. 2013 Jan 1;29(1):119–21.

Panconesi A, Sozio M. Algorithms in Bioinformatics. Springer; 2004. Fast hare: a fast heuristic for single individual SNP haplotype reconstruction; pp. 266?277.

Patterson M, Marschall T, Pisanti N, van Iersel L, Stougie L, Klau GW, Schnhuth A. WhatsHap: Weighted Haplotype Assembly for Future-Generation Sequencing Reads. J Comput Biol. 2015 Jun;22(6):498–509.

Pirola Y, Zaccaria S, Dondi R, Klau GW, Pisanti N, Bonizzoni P. HapCol: accurate and memory-efficient haplotype assembly from long reads. Bioinformatics. 2016 Jun 1;32(11):1610–7.

Tewhey R, Bansal V, Torkamani A, Topol EJ, Schork NJ. The importance of phase information for human genomics. Nat Rev Genet. 2011 Mar;12(3):215–23.

Wang Y, Feng E, Wang R. A clustering algorithm based on two distance functions for MEC model. Comput Biol Chem. 2007 Apr;31(2):148–50.

Weirather JL, de Cesare M, Wang Y, Piazza P, Sebastiano V, Wang XJ, Buck D, Au KF. Comprehensive comparison of Pacific Biosciences and Oxford Nanopore Technologies and their applications to transcriptome analysis. Version 2. F1000Res. 2017 Feb 3 [revised 2017 Jan 1];6:100.

Zook JM, Catoe D, McDaniel J, Vang L, Spies N, Sidow A, Weng Z, Liu Y, Mason CE, Alexander N, Henaff E, McIntyre AB, Chandramohan D, Chen F, Jaeger E, Moshrefi A, Pham K, Stedman W, Liang T, Saghbini M, Dzakula Z, Hastie A, Cao H, Deikus G, Schadt E, Sebra R, Bashir A, Truty RM, Chang CC, Gulbahce N, Zhao K, Ghosh S, Hyland F, Fu Y, Chaisson M, Xiao C, Trow J, Sherry ST, Zaranek AW, Ball M, Bobe J, Estep P, Church GM, Marks P, Kyriazopoulou-Panagiotopoulou S, Zheng GX, Schnall-Levin M, Ordonez HS, Mudivarti PA, Giorda K, Sheng Y, Rypdal KB, Salit M. Extensive sequencing of seven human genomes to characterize benchmark reference materials. Sci Data. 2016 Jun 7;3:160025.

